# Adhesion-driven vesicle translocation through membrane-covered pores

**DOI:** 10.1101/2024.05.20.594296

**Authors:** Nishant Baruah, Jiarul Midya, Gerhard Gompper, Anil Kumar Dasanna, Thorsten Auth

## Abstract

Translocation across barriers and through constrictions is a mechanism that is often used in vivo for transporting material between compartments. A specific example is apicomplexan parasites invading host cells through the tight junction that acts as a pore, and a similar barrier crossing is involved in drug delivery using lipid vesicles on the skin. Here, we use triangulated membranes and energy minimization to study the translocation of vesicles through pores with fixed radii. The vesicles bind to a lipid bilayer spanning the pore, and the adhesion-energy gain drives the translocation; the vesicle deformation while squeezing through leads to an energy barrier. In addition, the deformation-energy cost for deforming the pore-spanning membrane hinders translocation. Increasing the bending rigidity of the pore-spanning membrane and decreasing the pore size both increase the barrier height and shift the maximum to smaller translocation fractions. We compare the translocation of initially spherical vesicles with fixed membrane area and freely adjustable volume to that of initially prolate vesicles with fixed membrane area and volume. In the latter case, translocation can be entirely suppressed. Our predictions may help rationalize the invasion of apicomplexan parasites into host cells and design measures to combat the diseases they transmit.

## I. INTRODUCTION

In vivo, vesicles are abundant for transporting material within and between cells. Examples are synaptic vesicles in neurotransmission [1], extracellular vesicles that are involved in physiological processes and are proposed as drug-delivery vehicles [2], synthetic liposomes for drug delivery [3], and enveloped viruses, such as SARS-CoV-2 [4, 5]. Vesicles that deliver their content to host cells often fuse with the host’s plasma membrane and thereby directly deliver their material to the membrane and the cytosol [6, 7]. However, endocytosis of entire vesicles at the plasma membrane is also an important uptake mechanism for extracellular vesicles [8] and enveloped viruses [9]. Because mammalian cells usually feature a cortical cytoskeleton below their plasma membrane, vesicles that are being endocytosed may need to “squeeze” through the cytoskeletal network; for example, the typical mesh size of the spectrin cytoskeleton of human erythrocytes is 60 *−* 100 nm [10].

Adhesion, wrapping, and squeezing have also been hypothesized to be relevant for the entry of apicomplexan parasites into their parasitophorous vacuoles within the host cells [11–13]: both, Plasmodium and Toxoplasma, deform upon invading host cells and squeeze through a “tight junction” [14, 15], which appears as an electrondense zone in microscopy but whose architecture is not yet entirely understood. In Toxoplasma, an actin ring within the parasite is found at the constriction [14]. The deformation energy of an intact polymerized membrane, such as a cortical cytoskeleton, has a fixed connectivity of the polymers that suppresses a complete engulfment of particles [16, 17]. Therefore, for successful invasion of Plasmodium into human erythrocytes, a local disassembly of the cortical spectrin cytoskeleton has been hypothesized [18]; the surrounding intact cytoskeleton might constrict the parasite during invasion.

The translocation of vesicles through pores has been studied using theory and computer simulations for various systems that differ mainly by the driving force for translocation and the membrane’s elastic properties. Many studies have been motivated by drug delivery in the skin where an osmotic-pressure difference drives the translocation [19]. For partial-translocated initially spherical vesicles, the membrane deformation energy increases compared to free vesicles, and the vesicle volume decreases [20]. For identical pore radii, a pore with a finite length increases the translocation-energy barrier compared to a pore with a vanishing length because of the increased compression of the vesicle [21]. An exponential decay of the translocation time with increasing driving force has been predicted for fluid vesicles, whereas a power law-dependence is expected for polymerized vesicles [21, 22]. A power-law dependence has also been reported for the critical strength of a homogeneous field driving pore translocation of fluid vesicles [23]. While all vesicle studies discussed above assume a fixed membrane area and a variable vesicle volume, vesicles with fixed volume and variable area– where the membrane stretching energy dominates–have also been studied [24].

Here, we study the translocation of vesicles through circular, membrane-covered pores, see Fig. 1. We compare our predictions for initially spherical vesicles with fixed membrane areas and variable volumes with those for prolate vesicles with fixed membrane areas and fixed volumes [25]. Within the pore, the vesicles adhere to the membrane covering the pore, providing an adhesion-energy gain that drives translocation. In addition, the deformation-energy costs for the pore-spanning membrane hinder translocation, leading to a maximum energy barrier when less than half of the vesicle area has translocated. Using a continuum membrane model, we calculate the energy profile for vesicle translocation, translocation states and times. For large pores, the energy barrier can be considerably lower for prolate vesicles than for initially spherical vesicles without a target volume, resulting in translocation times that are orders of magnitude shorter. However, for small pores, prolate vesicles with fixed area and volume may not translocate through the pore at all.

**FIG. 1:**
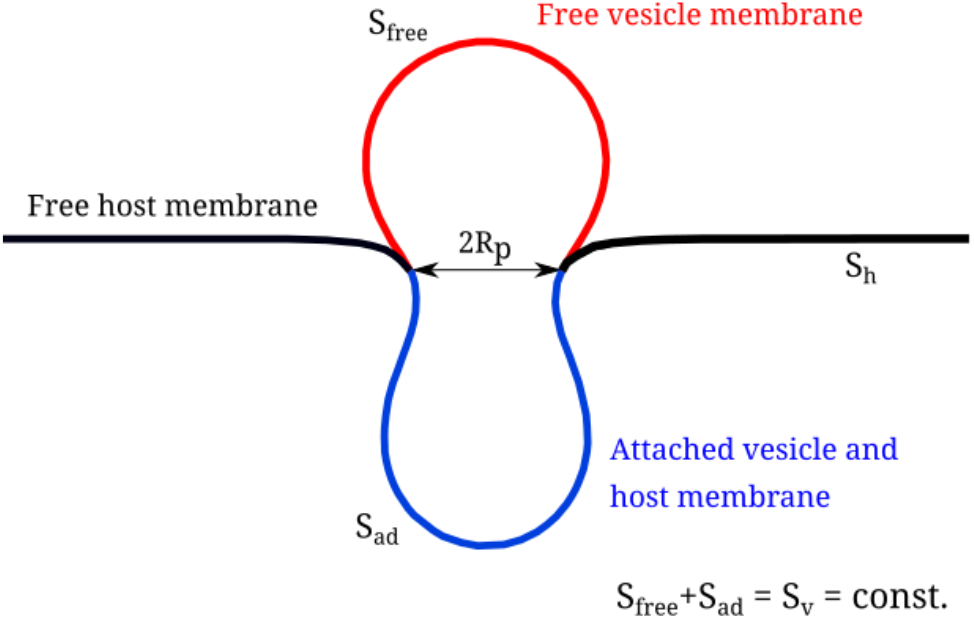
Initially spherical vesicle of radius *R*_v_ translocating through a circular pore of radius *R*_p_ = *R*_v_*/*2 in a host membrane with bending rigidity *κ*_h_ and tension 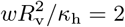. The snapshot corresponds to the vesicle-membrane translocation fraction *ρ* = 0.5; *S*_free_, *S*_ad_ and *S*_h_ indicate the free vesicle membrane, the double-bilayer of the vesicle membrane adhered to the host membrane that is bounded by the pore, and the free host membrane, respectively.

In section II, we introduce the model and present translocation-energy landscapes obtained using a spherical-cap geometry and triangulated membranes. In sections III and IV, we calculate stable translocation states, energy barriers, and translocation times for initially spherical and prolate vesicles, respectively. We characterize the dependence of the translocation transitions on membrane curvature-elastic parameters, pore and vesicle sizes, and vesicle adhesion strengths. Finally, in section V, we summarize our results and discuss their relevance for the invasion of deformable apicomplexan parasites into their host cells.

## II. MODEL AND METHODS

We study the translocation of a vesicle through a pore modeled by a circular ring of fixed radius embedded in a fluid membrane, see Fig. 1. In our calculations, the contact line between the vesicle and the pore-spanning membrane can have a wider radius than the pore; however, it coincides with the pore for all cases we have studied. The deformation-energy costs of the vesicle and the pore-spanning membrane hinder translocation, whereas the adhesion-energy gain for the contact between the vesicle and the pore-spanning membrane drives translocation. We discuss (i) initially spherical vesicles with fixed membrane area and freely adjustable volume and (ii) initially prolate vesicles with both a fixed area and volume. We calculate the translocation-energy contributions using a continuum membrane model,

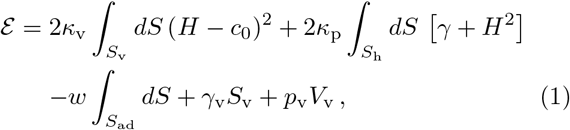

see Fig. 2. Here, *H* = (*c*_1_ + *c*_2_)*/*2 is the mean curvature, and *c*_1_ and *c*_2_ are the principal curvatures at each point of the membrane. The integrals are calculated over the membrane area *S*_v_ of the vesicle and the membrane area *S*_h_ of the pore-spanning membrane to that we refer in the following as host membrane. Furthermore, we refer to the area over which the vesicle membrane adheres to the host membrane as *S*_ad_ and to the free membrane area of the vesicle as *S*_free_. The membrane’s curvature-elastic properties are characterized by the bending rigidities *κ*_v_ of the vesicle and *κ*_h_ of the host; in addition, the vesicle membrane may be subject to a spontaneous curvature *c*_0_ and the host membrane to a tension *γ*. The vesicle-host contact interaction, which is applied within the pore only, is characterized by the adhesion strength *w*; the Lagrange multipliers *γ*_v_ and *p*_v_ fix the vesicle’s membrane area *S*_v_ and volume *V*_v_, respectively.

**FIG. 2:**
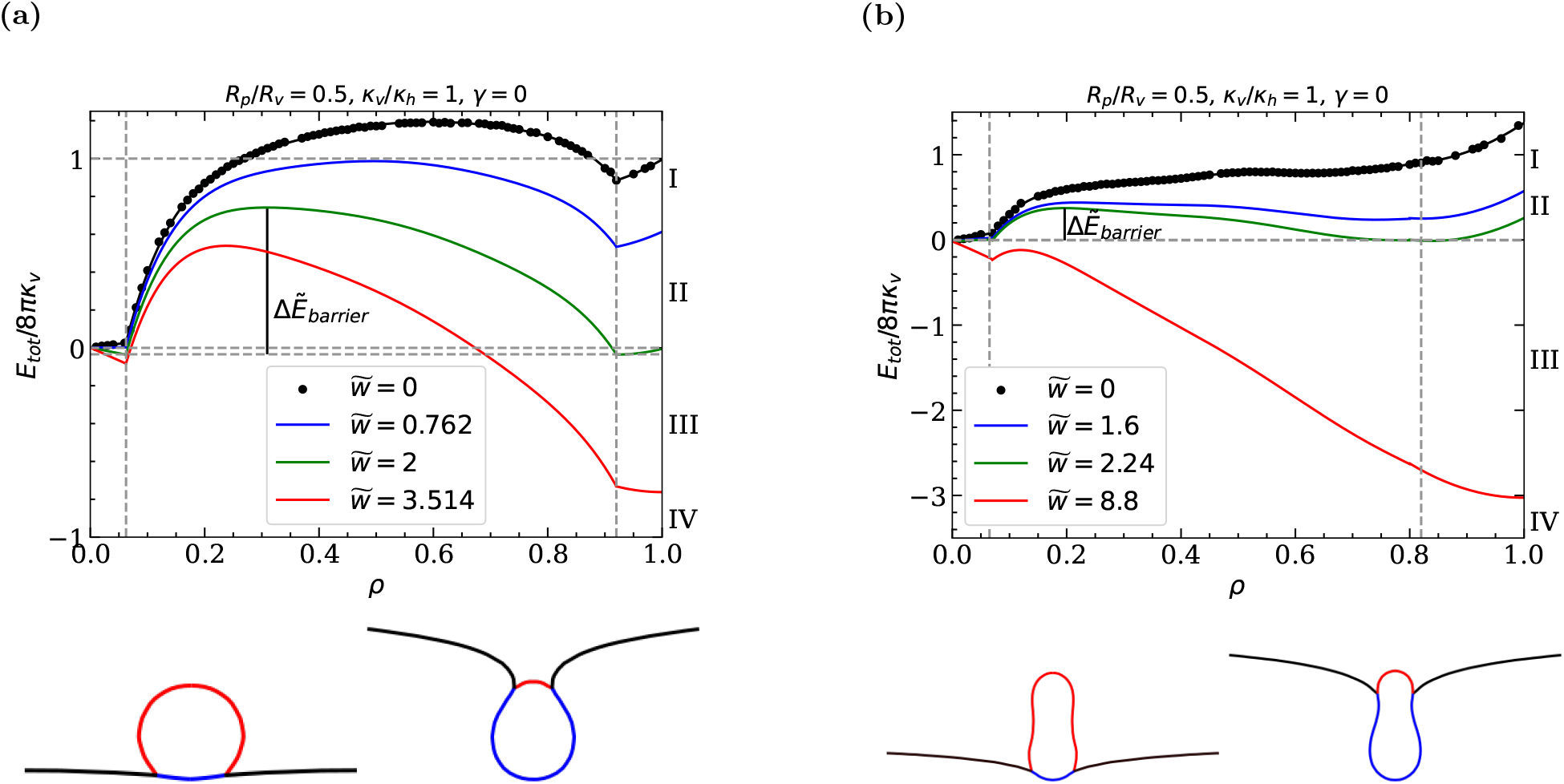
Energies and simulation snapshots for a vesicle-pore system with *R*_p_*/R*_v_ = 0.5 and *κ*_v_*/κ*_h_ = 1 as a function of the fraction *ρ* of translocated vesicle membrane area for (a) an initially spherical vesicle, and (b) a prolate vesicle with reduced volume *v* = 0.8. The points indicate the deformation energies calculated using triangulated membranes; the lines fits of piecewise functions to the data. The vertical dashed lines indicate the range of translocation fractions where the vesicle touches the rim of the pore. The total energies for reduced adhesion strengths 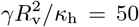 correspond to the transitions between free-vesicle (I), weak(II) and deep(III), and complete

The basis for all predictions of our model is the calculation of the total energy of the system for various fractions *ρ* = *S*_ad_*/*(*S*_free_ +*S*_ad_) of the vesicle’s membrane area having translocated through the pore, see Figs. 1 and 2. We first calculate the deformation energies only (*w* = 0) using either triangulated membranes [26, 27] and energy minimization to predict equilibrium shapes with the help of the freely available program “Surface Evolver” [28], or a spherical-cap model for that most calculations can be performed analytically, see Appendix A. For simplicity, we assume *c*_0_ = 0, although a finite spontaneous curvature affects particle wrapping by fluid membranes [29, 30] and may be induced in vivo, for example, by membranebound proteins [31, 32]. We characterize the vesicle size using the radius *R*_v_ of a sphere with the same surface area as the vesicle and the size of the circular pore using its radius *R*_p_. The prolate vesicles are, in addition, characterized by their reduced volume 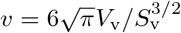 . -translocated states (IV). The reduced energy barrier for pore-passage transition II *→* III is 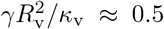 .

### A. Translocation-state transitions

The analysis of total energies for various adhesion strengths is performed analogously to the analysis for wrapping particles at membranes [33, 34]. Assuming a contact interaction between the vesicle and host membranes, the adhesion energy gain is proportional to the adhered membrane area and, thus, to the translocated fraction of the vesicle membrane, is added to the deformation energy costs. With the help of determining minima, maxima, and vanishing slopes in the deformation-energy landscape, transitions between stable translocation states, energy barriers, and translocation times are predicted. The triangulated-membrane data for the deformation energy is fit using a piecewise function *f* (*ρ*) that consists of fourth-order polynomials, whereas the deformation energies obtained using the spherical-cap model can be analyzed directly.

In general, we find the existence of four stable states in which the system can reside: in state I, the vesicle body has not contacted the host membrane; in state II, the vesicle is attached to the membrane, but most of its membrane area has not translocated through the pore; in state III, the majority of the vesicle membrane area has translocated, but the vesicle is not yet entirely enveloped by the host membrane; in state IV, the vesicle has translocated completely and is completely enveloped by the host membrane. We refer to the transitions I*→* II, II*→*III, and III*→*IV as binding, pore-passage, and envelopment transitions, respectively. In general, the translocation-energy landscapes for initially spherical and initially prolate vesicles differ significantly, see Fig. 2. However, for small translocation fractions *ρ*, the energy landscapes are qualitatively similar, and the deformation energy increases only weakly with increasing translocation fraction until the vesicle touches the pore. In this regime, the energies agree with those for the vesicles getting wrapped at membranes [35, 36].

As soon as the pore constricts the vesicle shape, the deformation energies increase steeply with increasing translocation fraction and experience a maximum. The energy barrier for the pore-passage transition is characterized by the value of *w* where the minima before and after pore-passage have equal energies, and the energy difference between the maximum and the minima, see Figs. 4 and 7, as well as the translocation fractions for energy minimums and maximums. For the same ratio of pore and vesicle radii, we find much lower energy barriers for the ‘thinner’ initially prolate vesicles than for initially spherical vesicles. In both cases, the maximum energy barrier for the pore-passage transition is found for less than half translocation. However, whereas the energy landscape shows a kink as a clear signature of detaching from the pore in the case of initially spherical vesicles, this feature is missing for initially prolate vesicles.

## III. TRANSLOCATION OF INITIALLY SPHERICAL VESICLES

Initially spherical vesicles can squeeze through arbitrarily small circular pores. However, unless the pore size is similar to the vesicle size, a typical energy barrier from the weak- to deep-translocated state has a height of the order of 8*π*(*κ*_v_ + *κ*_h_), see Fig. 2(a). Figure 3 shows translocation-state diagrams for various pore-to-vesicle size ratios, vesicle-to-host membrane bending rigidity ratios, and host membrane tensions. The state boundaries predicted using triangulated membranes are compared with estimations obtained using the spherical-cap model introduced in Appendix A.

**FIG. 3:**
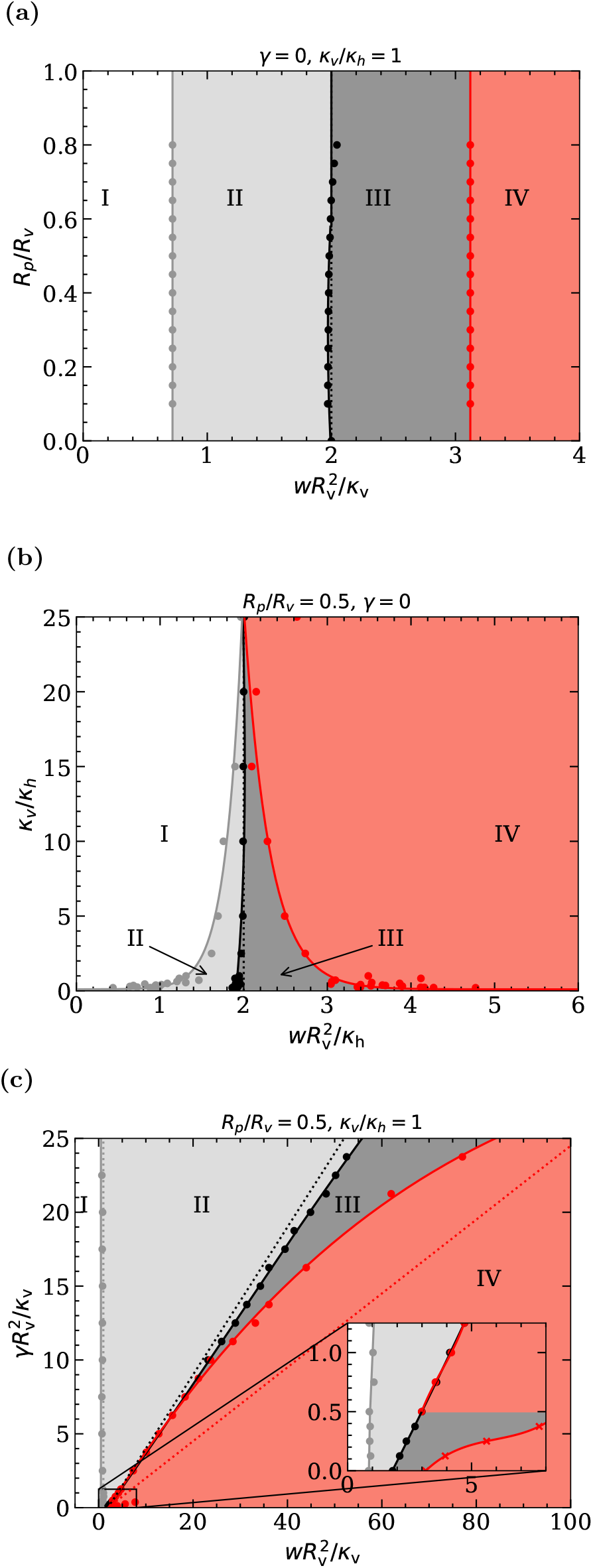
Translocation-state diagrams for initially spherical vesicles for (a) fixed *κ*_v_*/κ*_h_ = 1, various *R*_p_*/R*_v_ and adhesion strengths 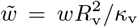 (b) fixed *R*_p_*/R*_v_, various *κ*_v_*/κ*_h_ and adhesion strengths 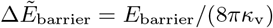 ; (c) fixed *R*_p_*/R*_v_, *κ*_v_*/κ*_h_ = 1, and various membrane tensions 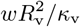 and adhesion strengths 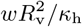 free-vesicle (I), weak-(II), deep-(III), and complete-translocated state (IV). Spherical cap model: dotted lines; Triangulated membranes: points and solid lines.

### A. Stable translocation states

Translocation-state diagrams for vesicles at tension-free host membranes show the existence of all four states for appropriate adhesion strengths, see Fig. 3(a,b). The vesicles do not adhere to the host membrane for small adhesion strengths. Assuming that the vesicles bind to the membrane in the center of the pore, the binding transition occurs at the adhesion strength known from particle-wrapping calculations. In this regime, for a hard sphere at a tensionless membrane, the deformation-energy profile is linear because the curvature of the particle surface is homogeneous [33, 34]. The vesicles flatten where they touch the host membrane, which increases the vesicle-host contact area and decreases the total energy for weak-translocated states [36]. Upon further translocation, the system assumes deep-translocated states and the complete-translocated state.

For fixed bending-rigidity ratio *κ*_v_*/κ*_h_, decreasing pore-to-vesicle size ratio *R*_p_*/R*_v_ decreases the adhesion strength for the transition between weak- and deep-translocated states, see Fig. 3(a). The transition is discontinuous with an energy barrier and thus corresponds to a jump between two stable states with different translocation fractions [37]. The adhesion strengths for the binding, pore-passage, and envelopment transitions are independent of *R*_p_*/R*_v_ and coincide with the predictions using the spherical-cap model.

For fixed pore-to-vesicle size ratio and high vesicle-to-host membrane bending rigidity ratios *κ*_v_*/κ*_h_, we recover the direct transition between the free and the complete-translocated state known for spherical hard particles, see Fig. 3(b). However, for small *κ*_v_*/κ*_h_ partial-translocated vesicles strongly deform, thus the binding transition shifts to lower and the envelopment transition to higher adhesion strengths. In particular, for *κ*_v_*/κ*_h_*→* 0, the binding transition approaches *w→* 0, whereas the envelopment transition shifts to higher adhesion strengths.

For various host-membrane tensions *γ*, the continuous binding transition is independent of *γ*, see Fig. 3(c), in agreement with particle-wrapping calculations [33, 38]. The pore-passage and envelopment transitions significantly shift to higher adhesion strengths with increasing *γ*. Interestingly, we find stable deep-translocated states below a threshold tension 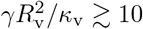 . For higher tensions, we observe a direct and discontinuous transition between the shallow-translocated and the complete-translocated state–with a metastable deep-translocated state. Finally, deep-translocated states are again stable for 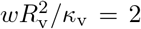, and the envelopment transition is continuous. The spherical-cap model predicts the binding transition at 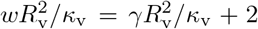 and the pore-passage transi-tion at 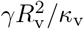 but cannot capture the envelopment transition.

### B. Energy barriers

Figures 4(a), (b), and (c) show the energy barriers for the pore-passage transitions for various pore-to-vesicle size ratios, vesicle-to-host membrane bending rigidity ratios, and host-membrane tensions, respectively, compare Fig. 2. In all cases, the energy barriers predicted using the triangulated-membrane model are lower than those estimated using the spherical-cap model. This is expected because the significantly higher number of degrees of freedom for the triangulated-membrane model compared with the spherical-cap model yields energetically more favorable vesicle shapes for partial-translocated states.

**FIG. 4:**
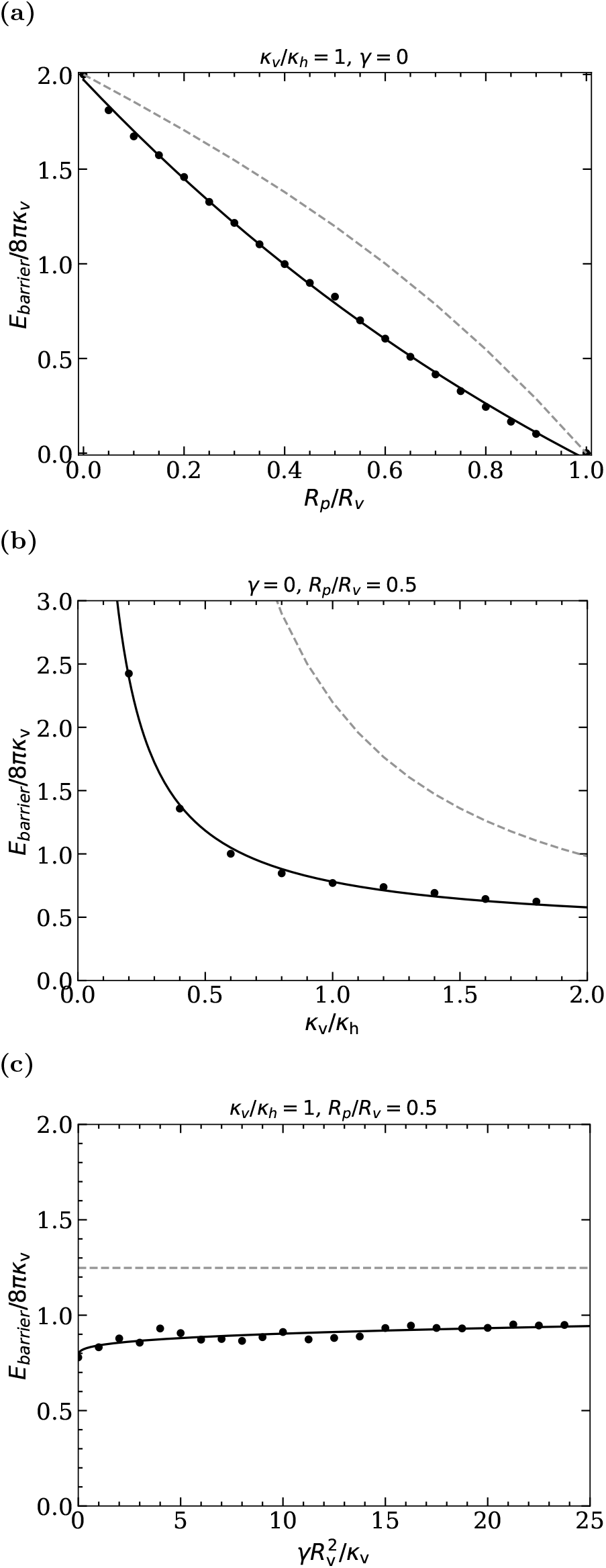
Energy barriers for the pore-passage transition of initially spherical vesicles and (a) *κ*_v_*/κ*_h_ = 1, *γ* = 0 and various vesicle-pore size ratios *R*_p_*/R*_v_ (b) *γ* = 0, *R*_p_*/R*_v_ = 0.5 and various vesicle-host bending-rigidity ratios *κ*_v_*/κ*_h_, and (c) *κ*_v_*/κ*_h_ = 1, *R*_p_*/R*_v_ = 0.5, and various host membrane tensions *γ*. Data is shown for the spherical-cap model (dashed lines) and triangulated membranes (points).

With increasing pore-to-vesicle size ratio *R*_p_*/R*_v_, the energy barrier decreases because the vesicle needs to deform less to squeeze through the pore, see Fig 4(a). The energy barrier vanishes for *R*_p_*/R*_v_ = 1, where the vesicle can translocate (almost) without being deformed by the pore. The maximal energy barrier for vanishing pore size is 8*π*(*κ*_v_ + *κ*_h_) for an infinitesimally small translocation fraction because the bending-energy costs jump from those for forming one to three complete vesicles. For small values *κ*_v_*/κ*_h_, the deformation of the host membrane contributes significantly to the energy barrier, whereas for high values of *κ*_v_*/κ*_h_ the barrier is dominated by the bending rigidity *κ*_v_ of the vesicle membrane both using the spherical-cap and the triangulated-membrane model, see Fig. 4(b). In the latter regime, the energy barrier is proportional to the bending rigidity *κ*_v_ of the vesicle only. With decreasing vesicle bending rigidity, the energy barrier will eventually depend on the host-membrane bending rigidity *κ*_h_ only.

Interestingly, the energy barrier increases weakly with increasing host membrane tension 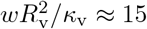 for the triangulated-membrane model and is independent of the tension for the spherical-cap model, see Fig. 4(c).

### C. Translocation times

We calculate translocation times using the Fokker-Planck equation for the ‘diffusion’ of vesicles across the energy barrier at finite vesicle-host adhesion strengths *w*.

For a Brownian particle subject to thermal motion, the translocation time is estimated based on the probability distributions for finding the particle at specific locations in the energy landscape [21],

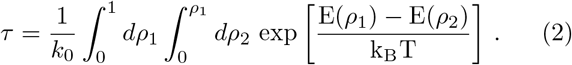

Here, *E*(*ρ*) refers to the difference of the total energy of the system calculated using Eq. (1) at translocation fractions *ρ*_1_ and *ρ*_2_, and *k*_0_ is a measure for the friction between the vesicle and the pore. Figure 5 shows the reduced translocation times *τ/τ*_0_ for initially spherical vesicles translocating through pores of various radii, where *τ*_0_ = 1*/k*_0_ [39].

**FIG. 5:**
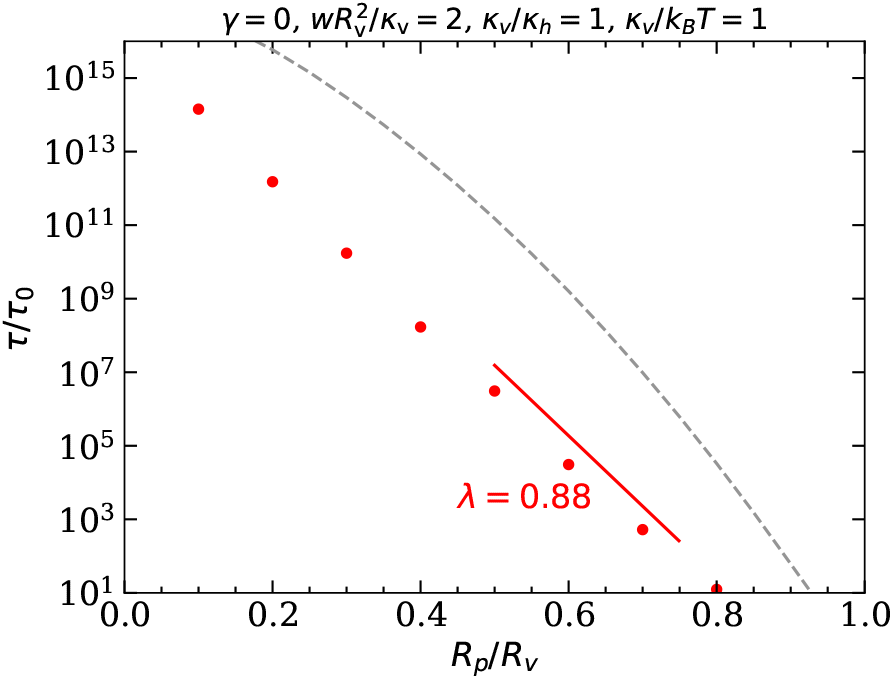
Translocation times for initially spherical vesicles with *R*_p_*/R*_v_ = 0.5, *κ*_v_*/κ*_h_ = 1, *κ*_h_*/k*_B_*T* = 1, and *γ* = 0 as a function of the pore-to-vesicle size ratio *R*_p_*/R*_v_, calculated using Eq. (2). Data is shown for the spherical-cap model (dashed lines) and triangulated membranes (points). The guide to the eye shows the characteristic exponential decay exponent *λ* in Eq. (3).

With the assumption of harmonic shapes for potential well and barrier, Eq. (2) can be reduced to the wellknown Kramer’s escape problem [40] with

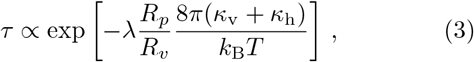

where the exponent is determined by the height of the energy barrier; we expect the proportionality factor to be of the order 1. For fixed adhesion strength, the translocation times decrease approximately exponentially with increasing pore-to-vesicle size ratio *R*_p_*/R*_v_, see Fig. 5. The exponential dependence of the translocation time on *R*_p_*/R*_v_ is expected from Eq. (3) because the barrier height *E*_barrier_ *≈* (1*− λR*_*p*_*/R*_*v*_)8*π*(*κ*_v_ + *κ*_h_) decreases almost linearly with increasing pore-to-vesicle size ratio, see Fig. 4(a). Thus, the translocation time shows a high sensitivity to *R*_p_*/R*_v_. The deviation of the exponential decay exponent *λ* from 1 accounts for the nonlinear dependence of the energy barrier on *R*_p_*/R*_v_, see Fig. 4(a), and the exact shape of the energy landscape.

In physiological conditions, we expect long translocation times because typical energy barriers are orders of magnitude higher than thermal energies. Furthermore, we find much shorter translocation times using triangulated membranes instead of the spherical-cap model, which reflects the lower energy barriers that we find for triangulated membranes and the importance of calculating accurate vesicle shapes.

## IV. TRANSLOCATION OF INITIALLY PROLATE VESICLES

The shapes of prolate vesicles that have a constant reduced volume, translocating through pores with radii comparable to the lengths of the vesicles’ short axes, are only weakly affected by the constriction. Thus, the energy barriers for prolate vesicles are smaller than those for initially spherical vesicles with the same membrane area, compare Fig. 2. However, complete translocation of prolate vesicles does not occur if the enclosed vesicle volume of half-translocated vesicles cannot be accommodated by the available vesicle membrane area, see Fig. 6(a). Furthermore, for vesicles with reduced volume *v* = 0.8, the range of adhesion strengths for stable deep-translocated states is considerably larger compared with initially the case of initiallt spherical vesicles. In this section, we discuss energy barriers and translocation times for cylindrically symmetric prolate-vesicle systems with reduced vesicle volumes *v* = 0.8 and *v* = 0.6 [41].

**FIG. 6:**
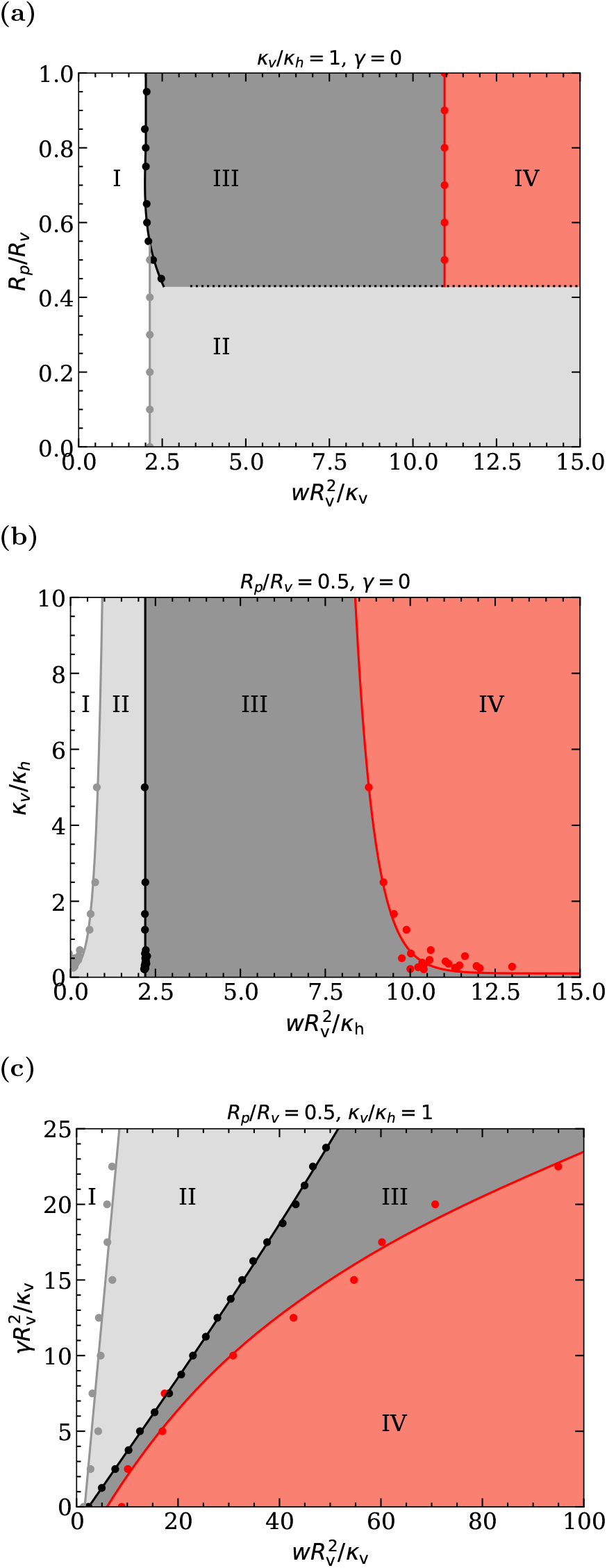
Translocation-state diagrams for prolate vesicles of *v* = 0.8, (a) fixed *κ*_v_*/κ*_h_ = 1, various size ratios *R*_p_*/R*_v_ and adhesion strengths 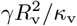 (b) fixed *R*_p_*/R*_v_, vari-ous bending-rigidity ratios *κ*_v_*/κ*_h_ and adhesion strengths 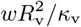 (c) fixed *R*_p_*/R*_v_, *κ*_v_*/κ*_h_ = 1, various membrane tensions 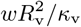 and adhesion strengths 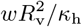 The states are labeled following Fig. 3.

### A. Stable translocation states

The translocation-state diagrams for prolate vesicles in rocket orientation with reduced volume *v* = 0.8 are qualitatively different compared with those for initially spherical vesicles, see, in particular, Figs. 6(a) and 3(a). The adhesion strengths for the binding transition are independent of the pore-to-vesicle size ratio and higher than for initially spherical vesicles because the highly curved tip of the vesicle has to be wrapped. For *R*_p_*/R*v ≳ 0.5, the binding transition is also the pore-passage transition and occurs directly from the free to the deep-translocated state; the deep-translocated state is stable over a large range of adhesion strengths. As for initially spherical vesicles, a continuous binding transition from the free to the shallow-translocated state occurs for *R*_p_*/R*v ≲ 0.45. However, the membrane area of the vesicle is too small to enclose the entire vesicle volume throughout the translocation process; the vesicle will remain shallow-translocated for all adhesion strengths above the binding transition. In-between, for 0.45 ≲ *R*_p_*/R*_v_ ≲ 0.5, we find transitions from the free to shallow-, deepand completewrapped states with increasing adhesion strengths. The range of adhesion strengths for stable shallow-wrapped states is small because of the high bending-energy costs for wrapping the highly curved vesicle tip. The continuous envelopment transition is independent of the poreto-vesicle size ratio.

We find a large region of stable shallowand deeptranslocated states in the translocation-state diagram for various vesicle-to-host membrane bending rigidities and adhesion strengths, see Fig. 6(b). The high deformation-energy costs for wrapping the highly curved tip of the vesicle to complete the translocation of a deep-translocated vesicle shifts the envelopment transition up to 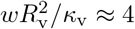, compared with 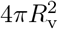 for initially spherical vesicles. For various host-membrane tensions, the adhesion strengths for the binding, pore-passage, and envelopment transitions increase with increasing host-membrane tension, see Fig. 6(c). Compared with initially spherical vesicles, the deep-translocated state is stable up to much higher adhesion strengths and exists for all host-membrane tensions.

### B. Energy barriers

The energy barriers for the translocation of initially prolate vesicles with reduced volumes *v* = 0.8 and *v* = 0.6 through pores with fixed radii are shown in Fig. 7(a), (b), and (c) with respect to the pore-to-vesicle size ratio, the bending-rigidity ratio of vesicle and host membranes, and the host membrane tension, respectively.

**FIG. 7:**
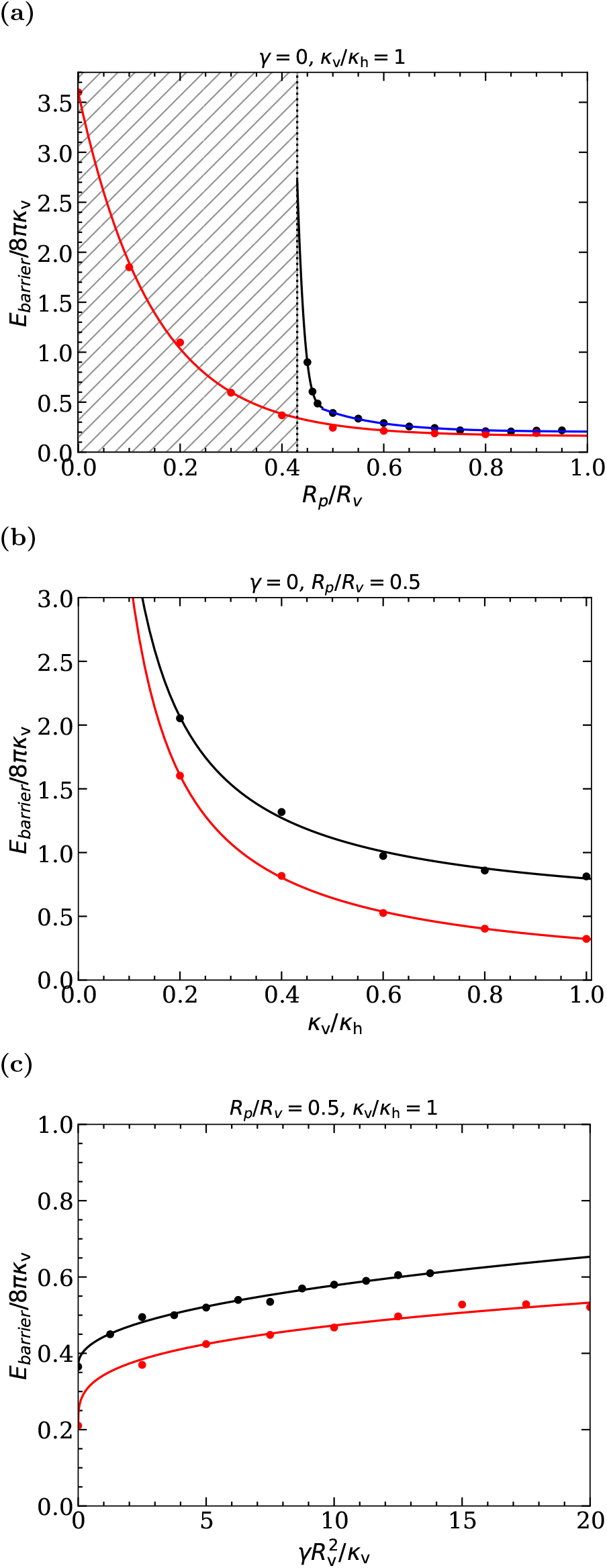
Energy barriers for the pore-passage transition of initially prolate vesicles with reduced volumes *v* = 0.6 (red) and *v* = 0.8 (black/blue), and (a) *κ*_v_*/κ*_h_ = 1 and *γ* = 0 as a function of *R*_p_*/R*_v_, (b) *R*_p_*/R*_v_ = 0.5 and *γ* = 0 as a function of *κ*_v_*/κ*_h_, and (c) *R*_p_*/R*_v_ = 0.5 and *κ*_v_*/κ*_h_ = 1 as a function of membrane tension 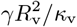 . In (a), the inaccessible parameter space for *v* = 0.8 is marked by the shaded region; the blue line indicates the barrier originating from wrapping the vesicle tip, and the black line the barrier originating from the pore constriction.

For pore-to-vesicle size ratios *R*_p_*/R*_v_ ≳ 0.6, the energy barrier for the pore-passage transition remains approximately constant because the vesicles are weakly or not at all constricted by the pore, see Fig. 7(a). The barrier originates from the high bending-energy costs for wrapping the tip of the vesicle and, therefore, does not vanish for *R*_p_*/R*_v_*→* 1. For *R*_p_*/R*_v_ ≲ 0.6, the dependence of the barrier height on the pore-to-vesicle size ratio is qualitatively different for the two reduced vesicle volumes. Whereas for *v* = 0.8 the energy barrier diverges at *R*_p_*/R*_v_*≈* 0.44 where the vesicle membrane area is too small to allow the vesicle to squeeze through the pore, for *v* = 0.6 the barrier increases smoothly with decreasing *R*_p_*/R*_v_ until an infinitesimally small pore size. The threshold reduced volume above that the energy diverges can be analytically calculated by equating the area for two equal-sized spherical caps connected to the pore for given available vesicle membrane area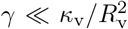,

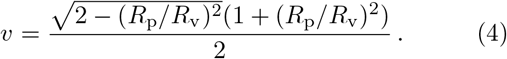

For varying bending rigidity ratio *κ*_v_*/κ*_h_ and *R*_p_*/R*_v_ = 0.5, the contribution of the host membrane dominates the energy barrier for small values of *κ*_v_*/κ*_h_ and has a negligible contribution for high values of *κ*_v_*/κ*_h_, see Fig. 7(b). For sufficiently large *κ*_v_*/κ*_h_ and *v* = 0.8, the energy barrier increases linearly with *κ*_v_, whereas for *v* = 0.6 the energy barrier vanishes (not shown). In the latter case, the vesicle translocates without being constricted by the pore, and the characteristic energy for deforming the host membrane is small compared with the characteristic energy for deforming the vesicle.

For *R*_p_*/R*_v_ = 0.5 and very small host-membrane tensions, 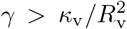, the energy barrier initially increases strongly with increasing *γ*, see Fig. 7(c). For large tensions, 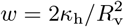 the energy barrier increases weakly with increasing host-membrane tension. The dependence of the barrier height on the host-membrane tension is qualitatively similar for both reduced volumes, but the absolute barrier height is lower for prolate vesicles with *v* = 0.6 compared to prolate vesicles with *v* = 0.8.

### C. Translocation times

The translocation times calculated using Eq. (2) show an exponential decrease with increasing pore-to-vesicle size ratio *R*_p_*/R*_v_, see Fig. 8. The decay constant is smaller for the initially spherical than for the prolate vesicles with *v* = 0.6, and typical translocation times are orders of magnitude longer for the initially spherical vesicles. In the case of the diverging energy barrier for *v* = 0.8, the translocation time also diverges. For high values of *R*_p_*/R*_v_, the translocation times for the prolate vesicles are independent of *R*_p_*/R*_v_, as are the energy barriers, see Fig. 7(a). However, the characteristic times for overcoming the wrapping-energy barrier for the tips are significantly smaller than those for the barriers originating from the pores constricting the vesicles.

**FIG. 8:**
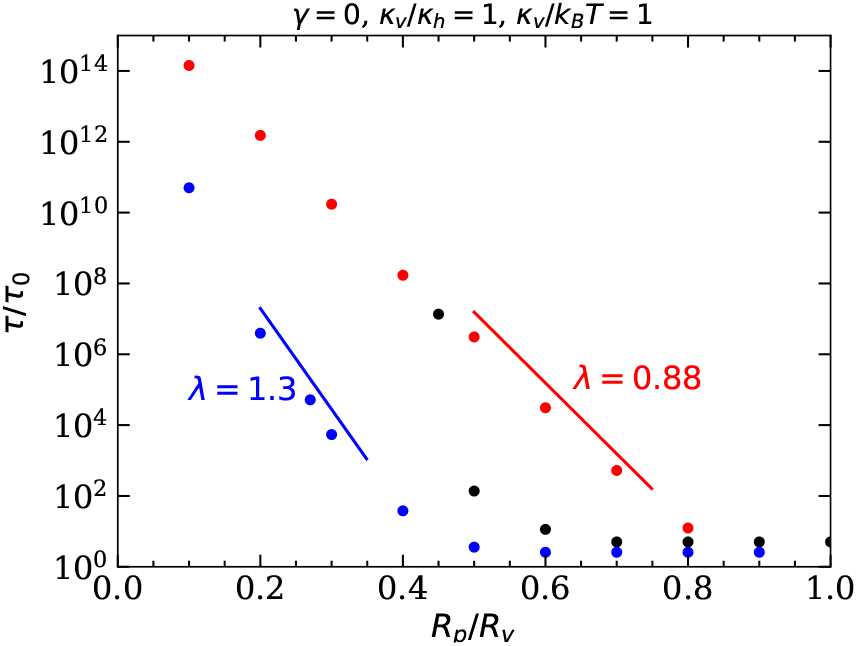
Translocation times for prolate and initially spherical vesicles with *κ*_v_*/κ*_h_ = 1, *κ*_h_*/k*_B_*T* = 1, and *γ* = 0 as a function of the pore-to-vesicle size ratio *R*_p_*/R*_v_, calculated using Eq. (2). Data is shown for an initially spherical vesicle (red), a prolate vesicle with *v* = 0.8 (black), and a prolate vesicle with *v* = 0.6 (blue). The guides to the eye show the characteristic decay exponents *λ* in Eq. (3).

## V. DISCUSSION AND CONCLUSIONS

In experiments, vesicle and cell shapes can often be quantified well using optical microscopy, but forces cannot be measured as easily. Computer simulations help us to connect vesicle shapes with energies and forces. Using a continuum-membrane model and exploiting cylindrical symmetry, we studied the translocation of initially spherical and prolate vesicles through circular pores with fixed radii. Vesicle translocation is driven by an adhesion-energy gain through a contact interaction with a pore-spanning host membrane, which at the same time contributes to the deformation-energy costs. The driving mechanism of adhesion is qualitatively different from translocation driven by an osmotic-pressure difference on both sides of the pore studied previously [20–22, 24]. Our results may help to understand cellular uptake processes, such as passive endocytosis of extracellular vesicles, budding of virions in the presence of a cortical host cytoskeleton, and invasion of parasites into host cells.

We find an exponential decay of the translocation times with increasing pore-to-vesicle size ratios. The deformation-energy landscapes have their maximum before half translocation because of the energy for deforming the host membrane. Significant translocation rates driven by thermal motion are expected for *κ*_v_*≈ κ*_h_ *≈ k*_B_*T*, for that the energy barriers are comparable to those for entropy-dominated systems, such as pore-passage of linear polymer chains [42–46]. Energy barriers for the polymers originate from the free-energy changes due to the reduced number of possible chain conformations and are therefore comparable to thermal energies. However, the barrier heights for vesicle-pore translocation are determined by the curvature-elastic properties of the fluid membranes, and physiological lipid-bilayer bending rigidities are of the order of *κ* = 50 *k*_B_*T* [47]. Therefore, a sufficiently high vesicle-host membrane adhesion strength is required that reduces the activation energy for experimentally observing translocation.

The established approach to simulate Plasmodium falciparum merozoites in the blood stage, based on electron-tomography data for free parasites, is hard egg-shaped particles [12]. For the reorientation and invasion of the parasites with their apical end towards the host cell membrane, mechanisms based on a receptor gradient on the parasite surface with a higher adhesion strength at the apical compared with the dorsal end [12] and on an interplay of red-blood-cell deformability and receptorligand bond dynamics [48, 49] have been hypothesized and quantified. However, recent microscopy images show that Plasmodium merozoites contain only two to three very short subpellicular microtubules [50] and experience major shape deformations during invasion [15, 51]. Furthermore, contrary to the widely accepted notion of invasion occurring from the merozoite’s pointed end, recent experiments indicate that invasion for Plasmodium knowlesi starts from the flattened end [52, 53]. Therefore, our results for vesicle-pore translocation provide a basis for studying the invasion of Plasmodium merozoites into red blood cells.

Our model of vesicles and fixed pores may be considered a biophysical approach for studying the host invasion of apicomplexans that captures the parasites’ prolate shape and finite deformability. However, for a pore with fixed radius, not including a cytoskeleton, and vanishing osmotic concentrations we may miss crucial mechanical aspects. In future, we plan to include further mechanical building blocks for modeling host entry of Plasmodium merozoites and Toxoplasma tachyzoites and to add active motor forces for invasion to the model.

## ACKNOWLEDGMENTS

NB, AKD, and TA acknowledge funding by the DFG SPP 2332 “Physics of Parasitism”, and helpful discussions with Mirko Singer (Heidelberg), and Javier Periz and Markus Meissner (Munich).

## Appendix A: Spherical-cap model

The energetics for an initially spherical vesicle translocating through a pore can be estimated using the spherical-cap model, where the fractions of the vesicle membrane on both sides of the pore are represented as spherical caps. Assuming vanishing spontaneous curvature, the integrals in Eq. (1) can be solved analytically,

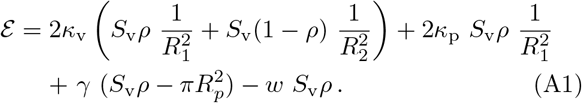

Here,

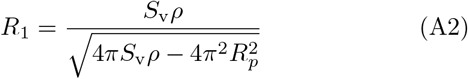

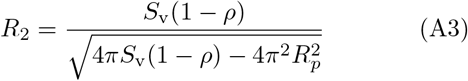

are the radii of the spherical caps for the membrane that has and has not translocated through the pore, respectively.

Equation (A1) is applicable when the vesicle touches the rim of the pore. When the vesicle does not touch the rim, in the spherical-cap model, it remains spherical with radius *R*_v_, and the total energy is the energy for wrapping a spherical particle,

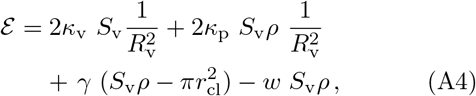

where *r*_cl_ *< R*_p_ is the radius of the contact line,

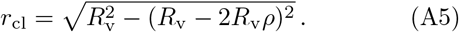

